# Metabolic modelling and time-resolved mapping of glucose oxidative metabolism in rats brain by indirect deuterium detection with ^1^H-FID-MRSI at 9.4T

**DOI:** 10.64898/2025.12.05.692506

**Authors:** Alessio Siviglia, Gianna Nossa, Brayan Alves, Fabian Niess, Anna Duguid, Zenon Starčuk, Wolfgang Bogner, Bernhard Strasser, Cristina Cudalbu, Bernard Lanz

## Abstract

**Object:** The present study exploits newly developed dynamic indirect ^1^H-[^2^H]-FID-MRSI at 9.4T, combined with a dedicated metabolic model, to enable regional and quantitative characterization of glucose oxidative metabolism flux in the rat brain with minimal metabolic assumptions, by measuring both ^2^H-labelled Glx turnover and pool size along a controlled ^2^H-Glc infusion protocol.

**Materials and Methods:** Seven rats underwent dynamic 2D ^1^H-FID-MRSI during a 2-hour infusion of [6,6’-^2^H_2_] glucose. Consecutive 13-min acquisitions quantified Glx-C4 ^1^H-signal decay, converted to ^2^H-Glx concentrations using baseline metabolite pool sizes. A three-pool kinetic model including ^2^H-label loss was fitted to regional turnover curves to estimate oxidative flux (V_gt_) and pyruvate dilution (Kdil). Model performance and parameter robustness were finally assessed with Monte-Carlo simulations.

**Results:** *In vivo* ^2^H-Glx turnover showed a saturated exponential rise (∼60 min), with a labelling plateau higher in striatum (1.85 μmol/g) than hippocampus (1.55 μmol/g). Metabolic modelling provided region-specific oxidative fluxes: V_gt_ = 0.27 ± 0.07 μmol/g/min (hippocampus) and V_gt_ = 0.40 ± 0.06 μmol/g/min (striatum), with consistent Kdil across regions. Simulations confirmed a good model robustness in retrieving V_gt_ over a large range of experimental conditions.

**Discussion:** This work shows the appropriateness of indirect dynamic ^1^H-[^2^H]-FID-MRSI for quantitative metabolic flux mapping of cerebral glucose oxidative metabolism.

## Introduction

The brain requires a continuous high level of energy, primarily depending on glucose (Glc) as its main energy source [1]. Because of this, studying and understanding Glc metabolism is an essential requirement for characterizing many neurological processes and diseases [2,3]. Common methods used to explore the Glc metabolic pathway *in vivo* are ^18^F-fluorodeoxyglucose (^18^F-FDG) positron emission tomography (PET) and carbon-13 magnetic resonance spectroscopy (^13^C-MRS). However, while FDG-PET can provide information about total brain Glc uptake, it cannot distinguish between the oxidative and non-oxidative Glc metabolic pathways, limiting the ability to discern between normal and pathologic Glc metabolism [4,5]. On the other hand, ^13^C-MRS is able to differentiate between those, but has an intrinsically low sensitivity due to its isotope’s low nuclear gyromagnetic ratio, which limits it to single voxel measurements in large volumes [6].

Deuterium metabolic imaging (DMI) has emerged over the last years as a powerful MR technique to probe the metabolism of labelled tracers such as Glc in the brain, enabling the mapping of downstream neurotransmitter synthesis in the form of combined glutamate+glutamine (Glu+Gln, Glx), and non-oxidative lactate (Lac) production [7-11]. Thus, deuterium magnetic resonance spectroscopic imaging (^2^H-MRSI) has become a robust technique with promising potential for clinical applications in various brain metabolism pathologies such as brain tumours and epilepsy [7,10,12].

Despite the limitations of a low gyromagnetic ratio (∼6.5 lower than ^1^H) and low natural abundance (0.015%, with geographical variability [12,13]), recent studies have highlighted some significant advantages of the ^2^H technique in the direct detection modality [12]. A common example is the relatively short T_1_ relaxation times for the ^2^H resonances of interest, following integration of ^2^H isotope after administration of deuterated tracers [7]. This particular feature overcomes the intrinsic low sensitivity of ^2^H by rapid signal acquisitions and averaging, thus, enabling metabolic imaging. Furthermore, ^2^H-MRSI is relatively insensitive towards magnetic field inhomogeneities and provides simple and clear spectra, allowing the characterization of Glc, Glx, and Lac - thanks to the naturally present deuterated water HDO component, used as reference - following administration of labelled Glc [10,14].

A drawback of the direct ^2^H detection modality is the need for specialized hardware, as well as the limited number of deuterated compounds that are estimated, which renders the MRSI technique less suitable for clinical settings. However, the indirect measurement of deuterated metabolism via ^1^H-MRSI may overcome these challenges. This is possible since deuterium cannot be detected with ^1^H-MRSI, and thus the ^1^H that are exchanged with ^2^H will not contribute to the proton spectrum, lowering the signal while still allowing for the quantification of the full metabolic profile [15]. Such an approach would allow for quantification of both metabolic pools size and labelled concentration using the same internal reference (water ^1^H signal). This excluded the need for ^2^H natural abundance assumptions, at the price of dealing with water residuals and lipid contamination of metabolite spectra. A similar detection protocol has been established in human research [15-19], although clear turnover curves kinetics have not been reported, nor interpreted and modelled in terms of metabolic fluxes. In addition, indirect ^2^H-MRSI has yet been shown preclinically [20]. Finally, mapping of cerebral metabolic kinetics requires high detection sensitivity, as well as, sufficient spatial and temporal resolution, which can be particularly challenging in small volumes of acquisitions encountered in rat preclinical studies.

To tackle these challenges, and building on recent ^1^H-FID-MRSI Quantitative exchange label turnover (QELT) developments [16,17], the aim of the present work is therefore to implement indirect detection of ^2^H labelling through ^1^H-FID-MRSI [21] in rats. This will take advantage of the higher sensitivity of ^1^H detection and the high spatial and time resolution at ultra-high field (9.4T) coupled with the use of a cryogenic RF probe, in order to map the dynamics of Glc oxidative metabolism in terms of decaying metabolite ^1^H signals, under continuous infusion of deuterated Glc.

Furthermore, specialized metabolic modelling [22] is required to generate quantitative maps (µmol/g/min) of oxidative and non-oxidative cerebral glucose metabolism. Such a model should consider a specific criticality relative to modelling of ^2^H-labelling, i.e. the presence of ^2^H label loss phenomena due to exchange with water ^1^H, which is expected to be significant mainly at the glycolysis and TCA cycle stages [23]. We thus present here a new metabolic model designed for *in vivo* ^1^H-FID-MRSI QELT data, which provides both metabolite pool sizes and dynamic ^2^H-labelling under a controlled blood ^2^H-Glc input function. This can enable the characterization of regional metabolic rates in the rat brain. The model’s performance was further assessed through Monte-Carlo (MC) simulations to test its robustness under varying experimental conditions.

## Methods

### In vivo experiments

The 2D ^1^H-FID-MRSI for ^2^H-labelling indirect detection was performed on n = 7 rats (mean weight = 248 ± 8 g) for *in vivo* brain data acquisition. The rats under isoflurane anaesthesia (1.5–2.0% in 0.9 l/min 50% oxygen/50% air mix, respiration rate kept at 60–70 breaths/min) were placed in the vendor-made stereotaxic holder (Bruker) with the use of ears and bite bars. Body temperature was maintained at 37.7 ± 0.2 °C by a warm water hose-circulating system. Respiration and body temperature were monitored via a small-animal system (SA Instruments, New York, NY, USA). The animals underwent overnight fasting prior to each experiment. All animal experiments were conducted according to federal and local ethical guidelines, and the protocols were approved by the local Committee on Animal Experimentation for the Canton de Vaud, Switzerland (VD 3892).

### Infusion Protocol

The glucose infusion procedure was adapted from the protocol described by Henry et al. ([24]) and used for ^13^C-MRS preclinical experiments [22,25,26]. An exponentially decaying bolus of 99%-enriched [6,6’-^2^H_2_] Glc (Sigma-Aldrich; 20% m/v in saline solution) was administered intravenously over 5 min, via an infusion pump and femoral vein cannulation, based on the measured basal glycemia and rat weight and aiming at 70% plasma fractional enrichment (FE), then maintained with a continuous infusion of 70%-enriched [6,6’-^2^H_2_] Glc (Sigma-Aldrich; 20% m/v in saline solution) for 2 hours. The infusion rate, calculated from a nominal whole body glucose disposal rate of 33.2 mg/kg/min [27], was further adjusted based on plasma glucose concentration concomitantly measured from a femoral artery, targeting a blood glycemia of 19 μmol/g. Blood sampling was carried out every 20 min throughout the duration of the infusion. In this way, a step-like input function for the blood Glc FE evolution was achieved, as well as the saturation of Glc transport across the blood brain barrier (BBB) (Figure 1) [28].

**Figure 1:**
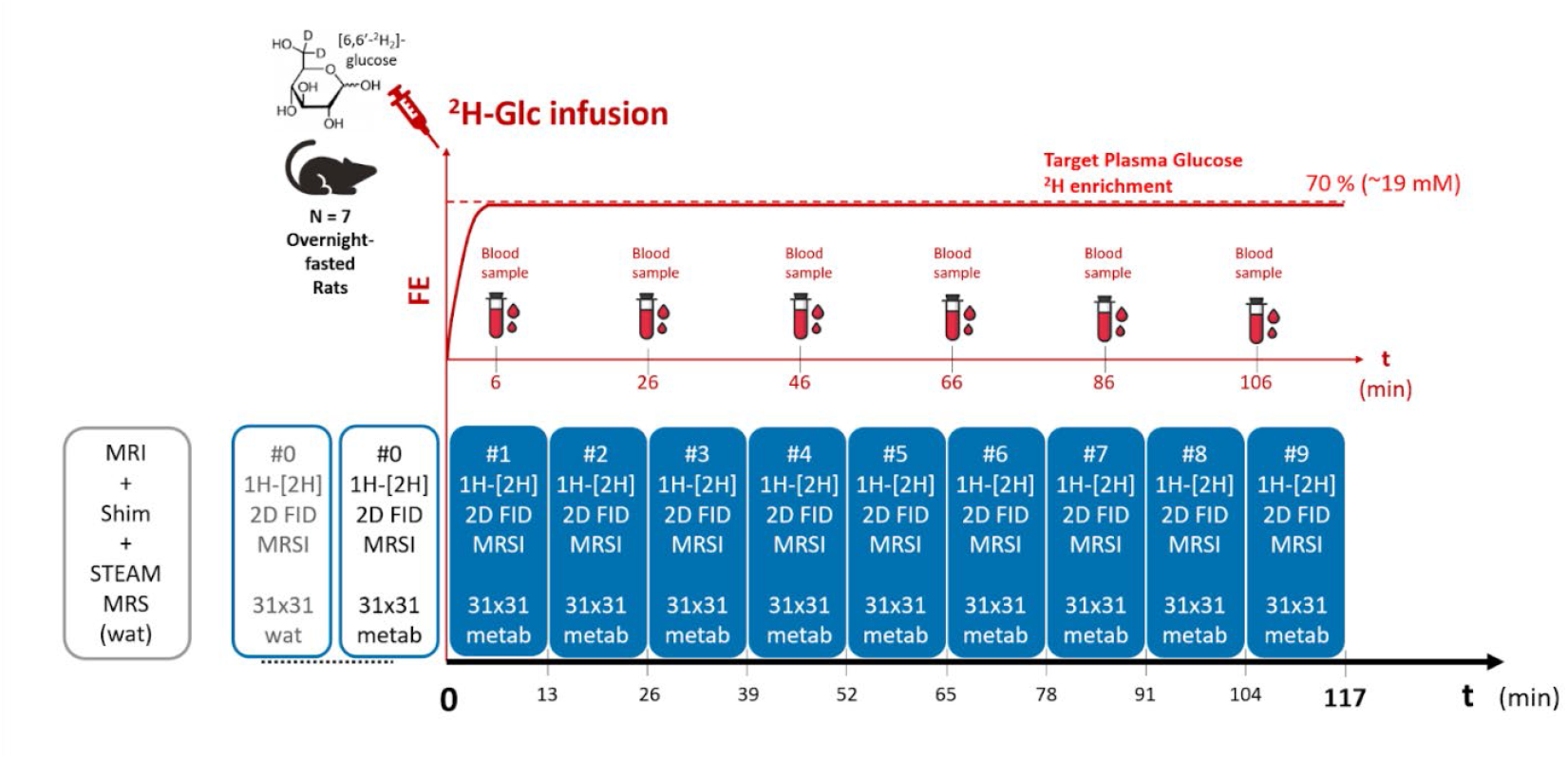
^1^H-[^2^H]-FID-MRSI protocol for indirect dynamic deuterium detection. Nine consecutive 2D ^1^H-FID-MRSI sequences (13 min) were performed during the ^2^H-Glc infusion, reaching an approximate time of 2h. The infusion rate was regulated based on arterial blood sampling every 20 min. A ^1^H-FID-MRSI acquisition (water + metabolites) was performed prior-infusion as a reference for the ^2^H-Glx concentration turnover quantification, as well as for Glx pool size characterization.

### Acquisition protocol

The experimental protocol was developed based on a 2D ^1^H-FID-MRSI acquisition scheme [29] adapted to a 9.4T system (Bruker/Magnex), using a volume transmit coil and receive-only 2 × 2 phased cryogenic array probe (CryoProbe, Bruker).

T2-weighted Turbo-RARE images were acquired in coronal and axial direction to position the MRSI slice and shimming volume, as well as for anatomical references (256 × 256 resolution, 20 slices, RARE_factor_ = 6, TR = 2500 ms, TE_eff_ = 33 ms, NA = 2, 0.8 mm slice thickness, FOV = 24 × 24 mm^2^). A second coronal T2-weighted Turbo-RARE image was acquired in each rat for brain segmentation (40 slices, TR = 4000 ms, TE_eff_ = 32 ms, NA = 10, RARE_factor_ = 6, 128 × 128 matrix, 0.3 mm slice thickness, FOV = 24 × 24 mm^2^).

The 2D FID-MRSI sequence consisted of a slice-selective Shinnar–Le Roux excitation pulse characterised by a bandwidth of 8.4 kHz, a pulse length of 0.5 ms, tuned to the Ernst angle of 55° for a TR of 823 ms and an acquisition delay (AD) of 1.3 ms including a 2D phase-encoding time of 0.5 ms. 768 spectral points were acquired with one average only, standard acquisition mode, and dealing with: a spectral bandwidth of 5 kHz (=dwell time of 200 μs), Cartesian k-space sampling, chemical shift offset of -2 ppm from the original working frequency of 4.7 ppm, 8 dummy scans, navigator (Gaussian 10° RF pulse, 1.37 ms duration, data size of 136, 23.87 ms module duration) and drift compensation. Hamming filtering was applied in the post-processing (Bruker Paravision 360 v3.5).

The 2 mm thick coronal MRSI slice was centered on the hippocampus and the striatum, with an acquisition matrix size of 31 × 31 over a FOV of 24 × 24 mm^2^ (Figure 2a), resulting in a nominal voxel size of 0.77 × 0.77 × 2 mm^3^ and in a total acquisition time of 13 min. The ^1^H-FID-MRSI sequence was combined with a VAPOR scheme and 12 FOV saturation slices for water and lipids signal suppression, respectively. First- and second-order shims were adjusted using Bruker MAPSHIM, first in an ellipsoid covering the full brain and further in a volume of interest (VOI) of 10 × 10 × 2 mm^3^ centered on the MRSI slice (Figure 2a). Shim quality was checked with a single-voxel STEAM spectroscopy water acquisition in the same VOI.

**Figure 2:**
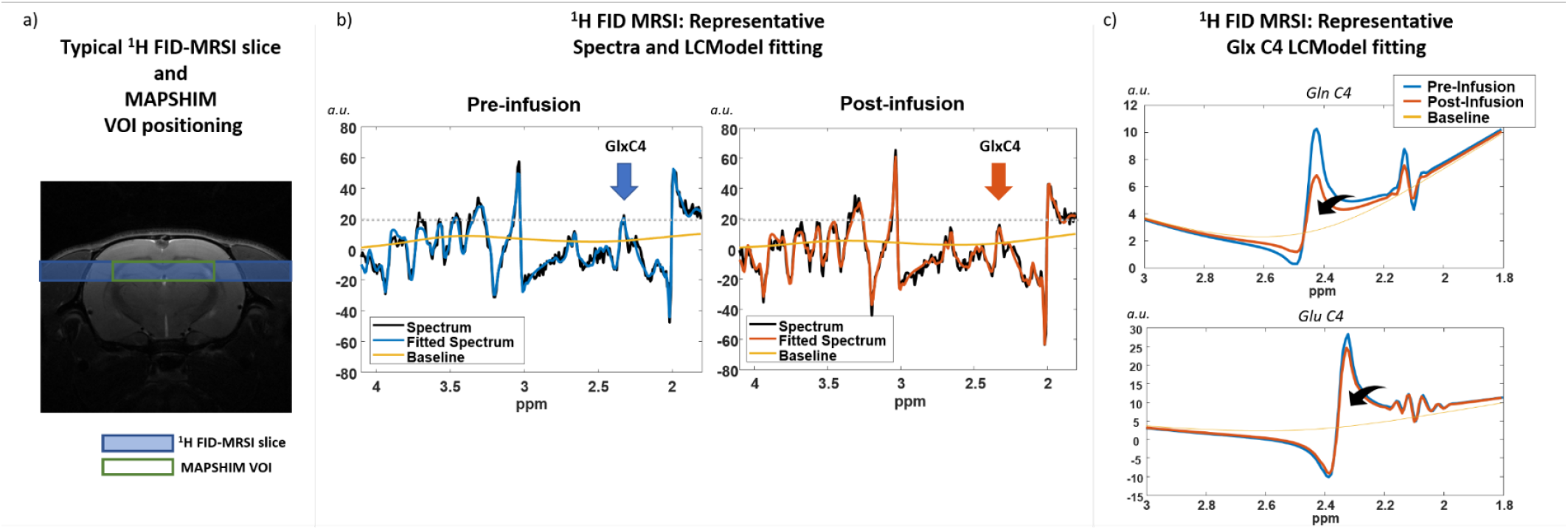
a) Representative ^1^H-FID-MRSI 2 mm slice positioning in the rat brain. Bruker MAPSHIM was applied in a 10 × 2 × 10 mm^3^ localized voxel overlapping with the slice, as well as a ^1^H STEAM water signal acquisition for shim quality check. b) Representative LCModel fitting of acquired ^1^H FID MRSI spectra, pre- and post-infusion (i.e. last infusion time-point). The highlighted Glx-C4 characteristic peak is expected to decrease along the infusion, as effectively detected by LCModel fitting of Glu and Gln spectral profiles (c).

After a baseline acquisition (Figure 1, iteration #0), the ^1^H-FID-MRSI sequence (acquisition time = 13 min, metabolites signal only) was repeated for the duration of the 2-hour ^2^H-Glc infusion protocol, for a total of 9 iterations.

### Data processing

Processing, quantifications, and metabolite maps were produced using the *MRS4Brain toolbox* with HLSVD water removal activated for MRSI [30], applied on k-space Hamming filtered data. Atlas-based segmentation using coronal T_2_-weighted acquisitions was applied for voxel selection in hippocampus and striatum. LCModel was used for metabolite fitting with a basis-set simulated in NMRScope-B/jMRUI [31-33], including ^2^H-labelled Glu and Gln patterns [16], and macromolecules signal acquired *in vivo* with the same FID-MRSI sequence. Quality control on both SNR (>4), mean linewidth (<1.25^*^mean) and CRLB (<40%) was applied. For all quantifications tCr was used as an internal reference set to 8 μmol/g.

The ^2^H-labelled Glx concentration obtained along the infusion was estimated by quantifying the decay of the ^1^H signal associated with the specific C4 position pattern of Glu and Gln. In particular, in the used basis-set both Glu and Gln profiles were made up of two separated components, i.e. the simulated C23 and C4 specific spectral patterns. In this way, for each ^1^H-FID-MRSI iteration along the infusion, the regional Glx-C4 relative decay with respect to the pre-infusion intensity was quantified via LCModel. Such a decay was then multiplied by the Glx pool size [Glx] measured by the baseline ^1^H-FID-MRSI acquisition, to obtain the time-point ^2^H-labelled Glx concentration of hippocampus and striatum.

### Metabolic modelling

The metabolic modelling work was based on previously validated ^13^C-labelling MRS approaches [22,25,26]. For the sake of simplicity, a unique astroglial compartment was considered for the description of ^2^H-labelled Glc metabolism, which includes both glial cells and neurons. The model was adapted from the biochemical pathways summarized in Figure 3a, consequent to the ^2^H-Glc ([6,6’-^2^H_2_] Glc) intake into the brain tissues and its metabolism, with a level of details adjusted to the amount of data accessible with *in vivo* ^2^H-MRS [23]. After the transport of Glc across the BBB and its phosphorylation, the deuterium atom is transferred from the C6 position across the phosphorylation process down to the C3 position of one pyruvate (Pyr) molecule after glycolysis. From the splitting process of a six-carbon molecule into two three-carbon molecules by the glycolytic chain, another unlabelled Pyr molecule is produced from each Glc molecule. From the Pyr pool, the ^2^H-label can be either channelled into the non-oxidative pathway to Lac through lactate dehydrogenase (LDH), or into oxidative metabolism in the TCA cycle, further reaching the C4 position of Glu within its first turn through transmitochondrial exchange from the TCA cycle intermediate 2-oxoglutarate [22]. Glu-C4 is further in exchange with Gln-C4 through the Glu-Gln cycle taking place in glutamatergic neurotransmission. Both pools were modelled as a combined kinetic pool Glx-C4 in this study. The main difference with ^13^C-label transfer is the partial ^2^H-label losses through exchange reactions with water hydrogens, in the glycolysis and the TCA cycle [12]. Further metabolism in the TCA cycle will lead to a complete label loss in the first TCA cycle turn.

**Figure 3:**
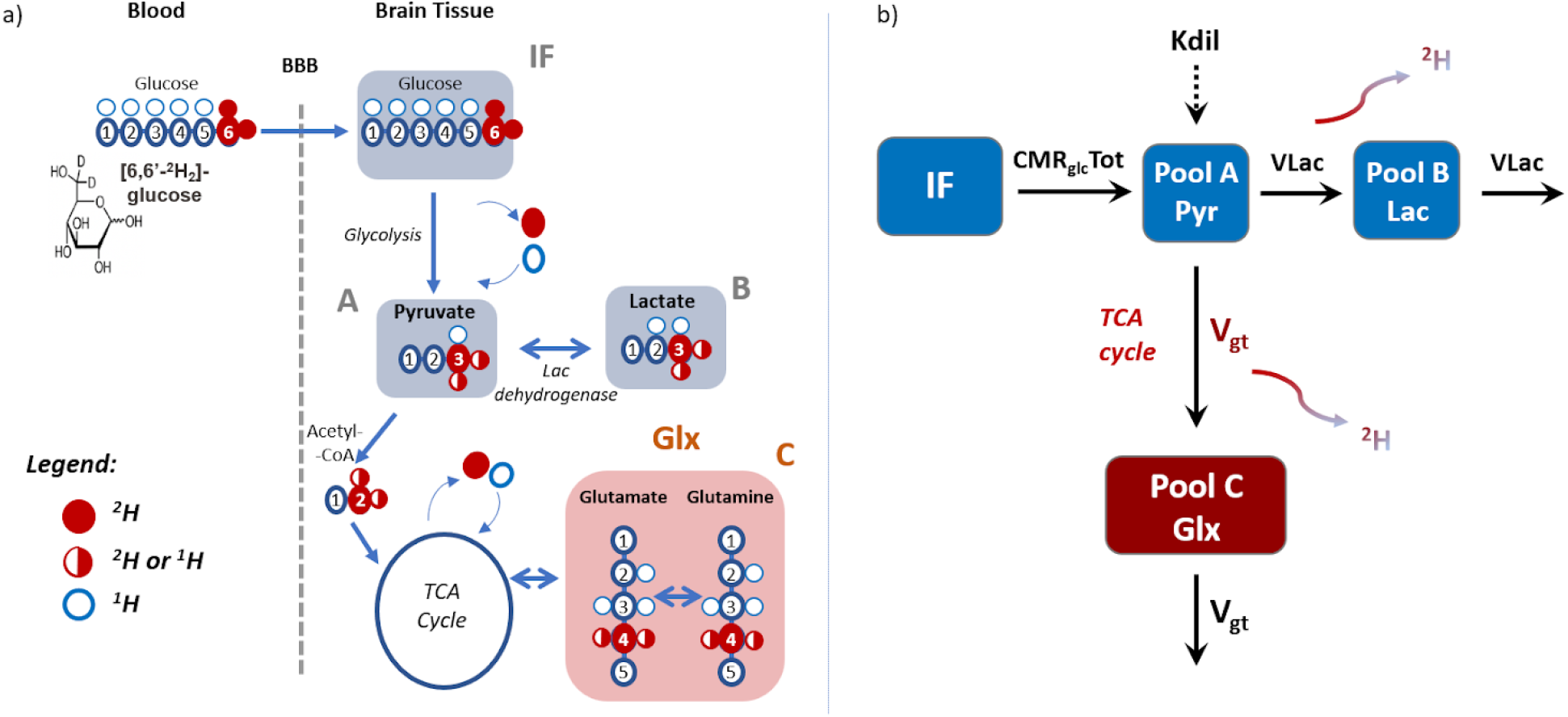
a) ^2^H-labelling scheme in the metabolism of [6,6’-^2^H_2_] Glc in brain tissues: after transport across the blood-brain barrier (BBB), ^2^H tracer reaches the position 3 of pyruvate through glycolysis from which it reaches the Glu and Gln position C4 through the TCA cycle in the oxidative pathway, or Lac C3 in the non-oxidative pathway through LDH. The ^2^H-label can be lost due to protium exchange with water mainly across the glycolysis and the TCA cycle pathways, and it is completely lost at the end of the first TCA cycle turn. b) The designed 3-pools metabolic model for ^2^H-Glc labelling experiments in brain tissue. The scheme models first the glycolysis process leading to the synthesis of pyruvate, from whose pool the two possible pathways are represented, i.e. LDH and TCA cycle. The cerebral metabolic rates characterising these two branches (V_Lac_ and V_gt_, respectively) and the pyruvate pool dilution factor taking into account Pyr production from other unlabelled substrates (K_dil_) are the free parameters of the model differential equations, solvable knowing the Glc input function, assuming a metabolic steady-state condition.

The designed kinetic model pools scheme (Figure 3b) consists of a first kinetic pool (A) for brain intracellular Pyr, from which two branches corresponding to the non-oxidative and oxidative energy metabolism connect to brain Lac (pool B) and the combined pool Glx (pool C). These branches respectively represent the Pyr-Lac exchange and the TCA cycle, characterized by the metabolic fluxes V_Lac_ and V_gt_. V_gt_ corresponds to the combined flux representing TCA cycle activity and transmitochondrial exchange with Glu [22]. The input function of the metabolic system is given by the glucose FE, represented by the transported ^2^H-Glc across the BBB, assumed to tightly follow the blood Glc enrichment, designed to reach a step function at 70% in less than 5 min. This is achieved by the saturation of the Glc transporters across the BBB with a maintained target glycemia around 19 μmol/g. Furthermore, ^2^H dilution processes resulting from the production of Pyr from alternative energy substrates were considered at the corresponding labelling pool (e.g. blood-borne lactate) with a global dilution factor Kdil (%). Assuming a metabolic steady-state condition, i.e. all the metabolic rates and total metabolites concentration assumed constant over time, and a comparable specific TCA cycle flux V_Tca_ to the TCA → Glx V_x_ transmitochondrial exchange flux (VTCA ≅ Vx, thus providing CMRglc,tot = (2 _Vgt+_V_Lac) / 2_ ) [25] [26] [34] [35], one can retrieve the following differential equations:

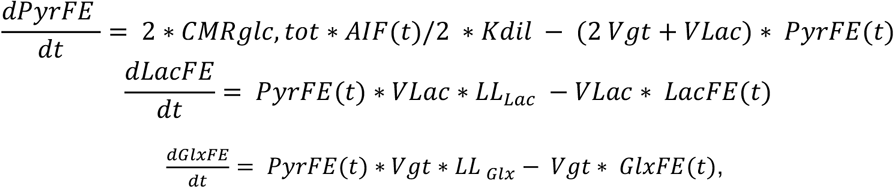

where LL_Lac_ and LL_Glx_ represent label loss factors (%) for Lac synthesis and from the TCA cycle to the Glx turnover, respectively [23].

The described model was implemented in Matlab (MathWorks, Natick, MA, USA). The model fitting to the obtained *in vivo* ^2^H-Glx turnover curves to retrieve V_gt_ and Kdil was performed via a standard built-in ordinary differential equation solver, combined with a modified Levenberg-Marquadt nonlinear regression method. The fitting was applied on the regional group-averaged curves, to retrieve the average rates for hippocampus and striatum over the cohort of n = 7 rats, as segmented from the MRI anatomical reference in the *MRS4Brain toolbox*.

### Metabolic modelling validation and robustness study

The model validation was carried out by performing n = 100 Monte-Carlo (MC) simulations on a scenario characterised by V_gt_ = 0.40 μmol/g/min and Kdil = 0.57 (estimations retrieved by the model application on *in vivo* striatum data), V_Lac_ = 0.20 μmol/g/min, Glc FE arterial input function AIF = 0.70 reached with a linear increasing ramp in 5 min, metabolite pool sizes [Pyr] = 0.1 μmol/g, [Lac] = 2.0 μmol/g, [Glx] = 14.0 μmol/g (estimation from *in vivo* striatum data), T_exp_ = 117 min and Δt=13 min to match the current experimental parameters for the ^1^H-[^2^H]-FID-MRSI protocol, nominal noise level for Glx **σ**_Glx_= 0.12 μmol/g (estimation by the fitting residuals on striatum data model application) and for Lac **σ**_Lac_ = 0.05 μmol/g as input values, accompanied by label loss factors retrieved from the literature (LL_Lac_ = 15.7% and LL_Glu+Gln_ = (37.9%+41.5%) / 2 [23]).

An extended MC test to assess the model robustness was performed to verify the accuracy and the precision of the model in retrieving the simulated flux rates V_gt_ = 0.40 μmol/g/min, V_Lac_ = 0.20 μmol/g/min and Kdil = 0.57 at the variation of experimental conditions as experiment time T_exp_, time resolution Δt and noise factor NF, defined as a multiplication factor of the nominal noise level. For each of these tests, default parameters values were T_exp_ = 117 min, Δt = 13 min, **σ**_Glx_ = 0.12 μmol/g/min and **σ**_Lac_ = 0.05 μmol/g/min, multiplied by a noise factor NF = 1. When investigated, T_exp_ was varied over a range of 60 -300 min, Δt over 2 - 30 min and NF over a range of 0.1 - 5.0, while keeping the other parameters constant. Both retrieved mean flux values and relative standard deviations were analysed. With an analogous approach, the model estimated parameters cross-correlation, i.e. the independent estimation of V_gt_ and V_Lac_, was also investigated by testing the stability of the retrieved nominal input flux value while varying the other flux value in the range of 0.05 - 1.00 μmol/g/min.

As for the metabolic modelling of the *in vivo* data, the model fitting to all the MC realisations was obtained with the same metabolic model via a standard built-in ordinary differential equation solver, with a modified Levenberg-Marquadt nonlinear regression method. Random initial conditions for the free parameters V_gt_, V_Lac_ and Kdil over a range of 0.01 - 1.00 were chosen for each MC realisation, to avoid systematic biases due to underexploration of the parameter space and minimizing the likelihood of ending in a local minimum of the regression [35].

## Results

### ^1^H-FID-MRSI protocol for indirect ^2^H-labelling detection

Plasma Glc quickly rose to levels around 19 μmol/g in less than 5 minutes and was maintained over the 2h infusion. LCModel fitting of pre- and post-infusion ^1^H-FID-MRSI acquired spectra showed a specific labelling pattern of the C4 position of Gln and Glu through ^1^H signal decrease (Figure 2b-c, 2.3-2.4 ppm), as expected following oxidative metabolism of [6,6’-^2^H_2_] Glc.

Pre- and post-Glc infusion ^1^H-FID-MRSI showed similar concentration levels for the Glx-C2 and C3 positions across the brain (average regional signal variations < 2%, Figure 4a,b,c), while the Glx-C4 hydrogen resonance showed a consistent decrease post-infusion (120 min) across the brain (Figure 4d,e,f), with -12% and -13% for Glx-C4 in the hippocampus and striatum, respectively (Figure 5b).

**Figure 4:**
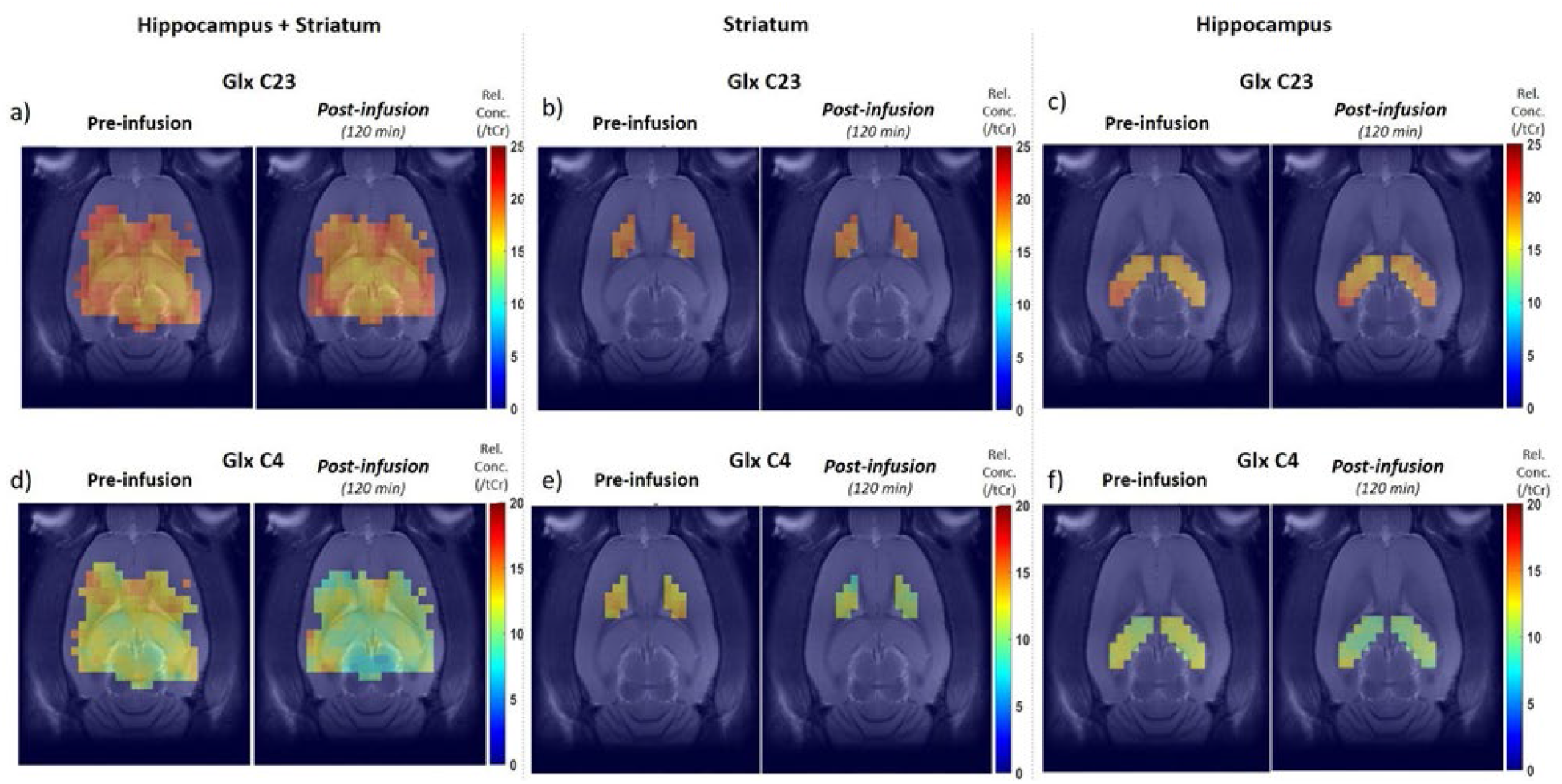
Representative metabolic concentration maps acquired in the rat brain. The maps show a steady concentration profile for Glx-C23 component over the brain (a), the striatum (b) and the hippocampus (c) along the infusion, while Glx-C4 decreases due to the progressive 2H turnover (d, e, f). The average Glx-C4 concentration decrease between the pre-infusion and 2h [6,6’-^2^H_2_] Glc infusion was -13% in the striatum and - 12% in the hippocampus. The quantifications were performed using tCr = 8 μmol/g as internal reference.

**Figure 5:**
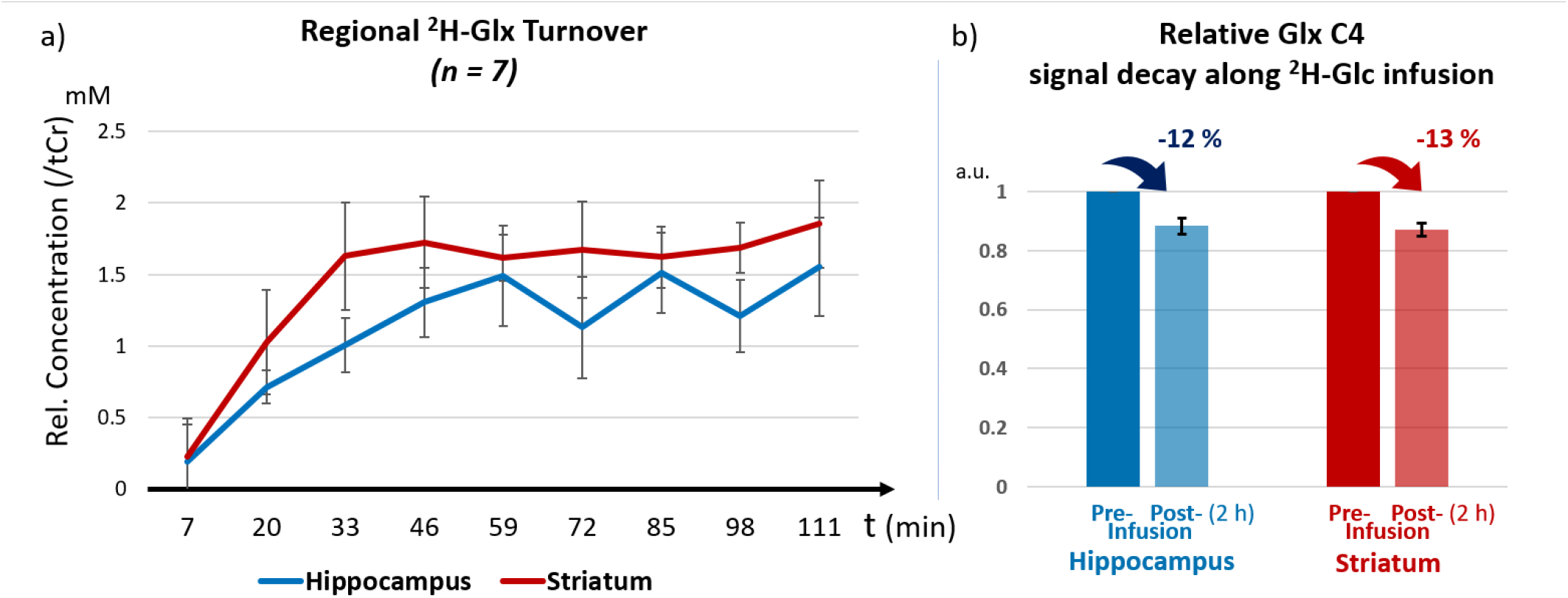
a) Group-averaged regional ^2^H-Glx concentration turnover curve along the ^2^H-Glc infusion (n = 7) in the rat hippocampus and striatum. Each time point is the quantification of a single 13 min ^1^H-FID-MRSI acquisition, using as total concentration reference the pre-infusion analogous acquisition result (in all cases tCr was used as internal reference, set at 8 μmol/g). b) Average relative Glx-C4 signal decay along ^2^H-Glc infusion, calculated on the last ^1^H-FID-MRSI iteration outcome with respect to the baseline (n = 7 rats), after 111 min infusion.

MRSI Glx-C4 signal decay was converted into ^2^H concentration by multiplying the percentage signal loss with the Glx pool size, as measured by ^1^H-FID-MRSI prior infusion. Figure 5a shows the resulting group-averaged ^2^H-Glx labelling concentration turnover for the hippocampus and the striatum, as measured with dynamic ^1^H-[^2^H]-FID-MRSI with a temporal resolution of 13 min. The curves show a rapid initial rise further reaching a labelling plateau after about 60 min, consistent across the animals. The Glx-C4 labelling curves also show consistent regional differences across the animal group between hippocampus and striatum, reaching a plateau concentration value for ^2^H-Glx of 1.55 μmol/g and 1.85 μmol/g, respectively. Furthermore, the curve initial rise appeared faster in the striatum (in about 40 min) than in the hippocampus (about 60 min).

### Metabolic modelling

The presented metabolic model successfully described the measured turnover of Glx-C4. Its fitting to the *in vivo* data (Figure 6) provided different outcomes for hippocampus and striatum data, *V*_*gt*_ = 0.267 ± 0.066 μmol/g/min with Kdil = 0.529 ± 0.030 and *V*_*gt*_ = 0.404 ± 0.055 μmol/g/min with Kdil = 0.566 ± 0.017, respectively.

**Figure 6:**
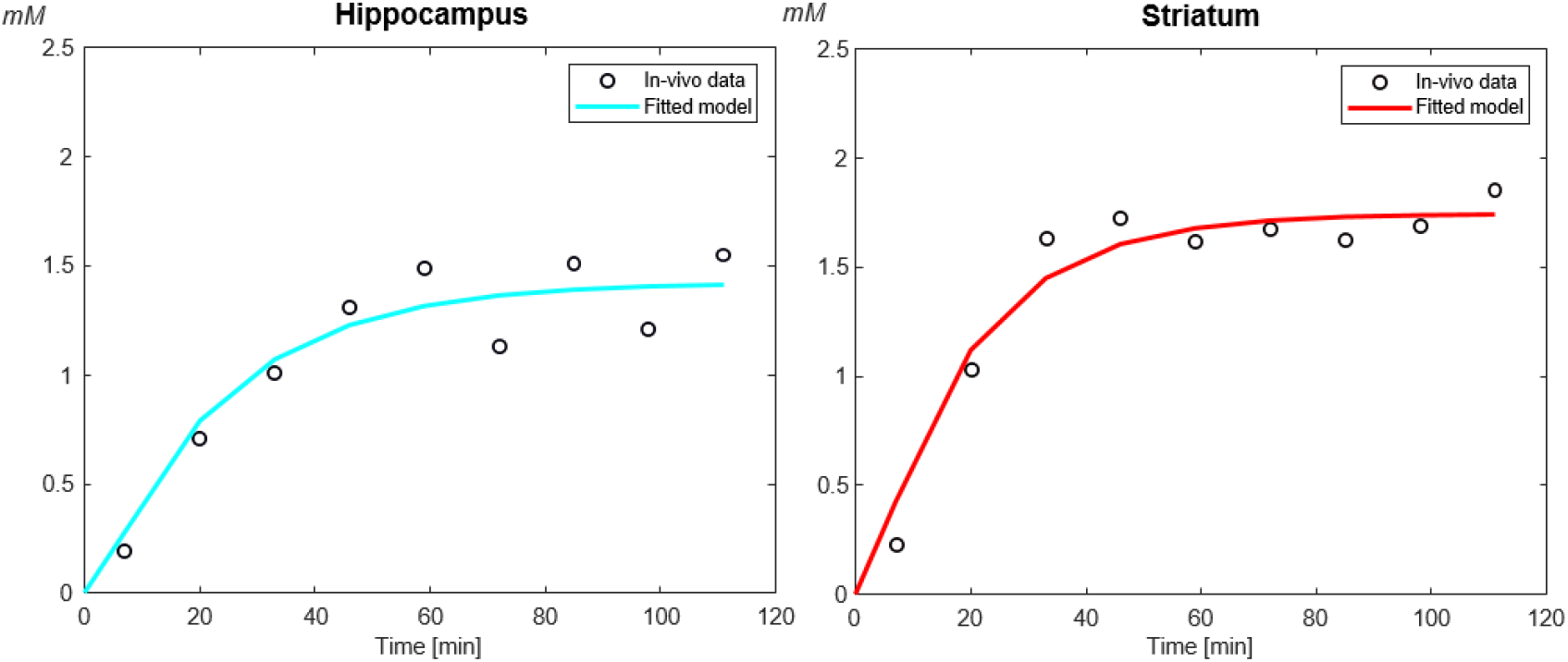
^2^H-Glx turnover along 2 hours infusion: in vivo data (group average, n = 7) vs fitted metabolic model for hippocampus (left) and striatum (right). The application of the metabolic model led to V_gt_ = 0.267 ± 0.066 μmol/g/min for hippocampus and V_gt_ = 0.404 ± 0.055 μmol/g/min for striatum.

Figure 7a shows the time-course for the labelled metabolites of interest as predicted by the designed metabolic model, in presence of a step-like AIF (^2^H-Glc FE plateau reached in 5 min), with pool sizes of 0.1 μmol/g, 2 μmol/g, and 14 μmol/g for Pyr, Lac, and Glx. The solutions consisted of typical exponential turnovers reaching plateau values of 0.02 μmol/g, 0.39 μmol/g and 2.65 μmol/g, approximately (with assumed input metabolic rates values of V_gt_ = 0.40 μmol/g/min and V_Lac_ = 0.20 μmol/g/min). Taking into account probabilities of label loss for Lac and Glx (LL_Lac_ = 15.7% and LL_Glu+Gln_ = (37.9% + 41.5%) / 2) [23]), the plateau values lowered to approximately 0.33 μmol/g and 1.59 μmol/g, respectively. Moreover, the turnover curves are characterised by different rising times, with around 80 min for labelled Glx and 20 min for labelled Lac to reach the plateau.

**Figure 7:**
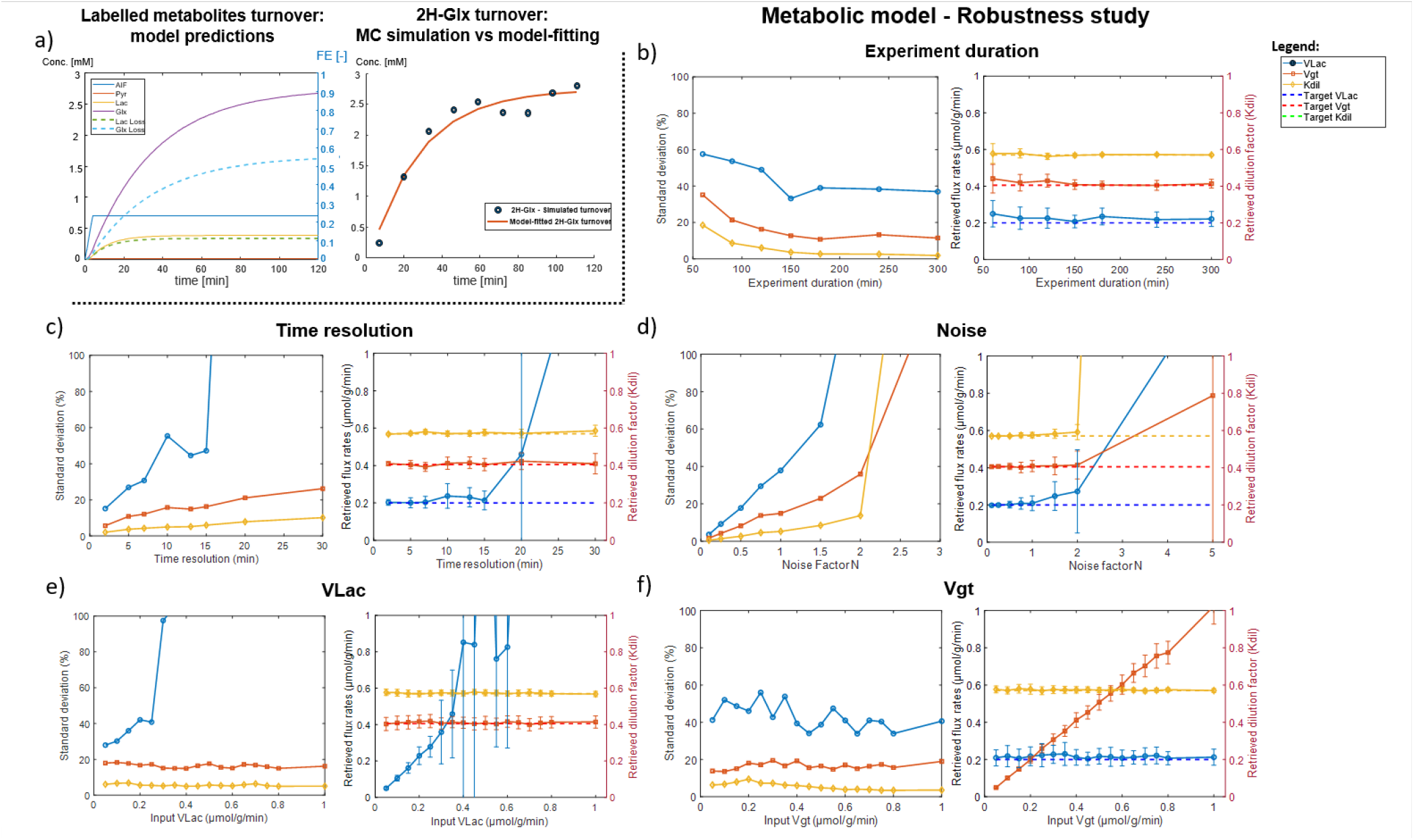
Metabolic model predictions (a) and robustness study (b-f). The labelled metabolites turnover curves (a) are predicted with V_gt_ = 0.40 μmol/g/min, V_Lac_ = 0.20 μmol/g/min, Kdil = 0.57 and a step-like AIF = 0.70. Both the model outcomes without (solid) and with (dashed) label losses (LL_Lac_ = 15.7% and LL_Glu+Gln_ = (37.9s% + 41.5%) / 2) are represented. The robustness study consisted of testing the precision (left plots, rel. standard deviations) and the accuracy (right plots, mean flux values) of the proposed model on N = 100 MC simulations varying the experiment duration (b), the temporal resolution (c), the data noise factor (d), V_Lac_ (e) and V_gt_ (f).

The presented metabolic model applied on N = 100 MC simulations (Figure 7a) of ^2^H-labelled Glx concentration evolution following a step-like input function for ^2^H-Glc provided an outcome of *V*_*gt*_ = 0.41 ± 0.06 μmol/g/min, *V*_*Lac*_ = 0.21 ± 0.10 μmol/g/min and Kdil = 0.57 ± 0.03, starting from pre-set values of *V*_*gt*_ = 0.40 μmol/g/min, *V*_*Lac*_ = 0.20 μmol/g/min and Kdil = 0.57.

The robustness study results (Figure 7b-f) showed a significant precision (relative standard deviations relStd < 20%) in retrieving V_gt_ in the different experimental parameter tests for, T_exp_ > 90 min, Δt < 20 min, NF < 1.5 and for the whole tested V_gt_ range (0.05 - 1.00 μmol/g/min). Furthermore, varying V_Lac_ over the same set of rate values resulted in a V_gt_ relStd < 20% for all the inputs, underlining an independence of the oxidative rate characterization from the non-oxidative flux. Regarding V_Lac_, its estimations precision is characterised by relStd < 20% for Δt < 5 min and NF ≤ 0.5 only, while a good accuracy (relative discrepancy by the input V_Lac_ rate < 20%) was retrieved for the tested T_exp_ > 60 min, Δt < 20 min, NF < 1.5 and V_Lac_ < 0.35 μmol/g/min ranges.

## Discussion

The present study shows for the first time the ability of ^1^H-FID-MRSI to detect quantitative signal changes at specific Glx molecular positions across a full rat brain slice over an extended 2h dynamic acquisition concomitant with the infusion of [6,6’-^2^H_2_] Glc. This enabled the characterization of Glx-C4 turnover with a sufficient temporal resolution to derive glucose metabolic fluxes. The significant signal variation (-12% for hippocampus and -13% for striatum) detected for the C4 position signal as opposed to the negligible change for C23s position (regional signal variations < 2%) underlined the reliability of the methodology to quantify indirectly the ^2^H-labelling in the rat brain. This was enforced by the high reproducibility of the measured turnover curves over the subject group (Figure 5a), obtained with a controlled [6,6’-^2^H_2_] Glc input function. The obtained typical exponential turnover curves and their turnover times were found to be in very good agreement with the labelling turnover of the same Glu and Gln positions achieved with ^13^C-Glc labelling experiments, reaching a labelling steady-state in about 60 - 80 min [25,36]. The measured concentration values were also coherent with the previous comparable ^13^C-Glc metabolic studies when considering the typical ^2^H label losses, as illustrated in the metabolic model prediction results (Figure 7a), reaching around 1.8 μmol/g ^2^H-Glx-C4 concentration for striatum (Figure 5a). This underlines the strong potential of the dynamic ^1^H-[^2^H]-FID-MRSI indirect detection approach to quantify metabolic rate maps.

A strong point for ^1^H-[^2^H]-FID-MRSI as compared to single voxel ^13^C-MRS was illustrated with its ability to detect regional metabolic differences in glucose oxidative metabolism reflected in the ^2^H-Glx turnover (Figure 5a,b): while for the striatum, a relatively rapid rise to reach the labelling steady-state (about 45 min) was observed, for the hippocampus, the rise was slower (about 60 min) and the ^2^H concentration plateau was lower (1.55 μmol/g compared to 1.85 μmol/g, respectively). The difference in the labelling rising phase suggests a slower V_gt_ in the hippocampus, while the lower plateau suggests a stronger dilution related to metabolism of alternative substrates in this region. This was reflected in the metabolic modelling of the ^2^H turnover curves with the adapted compartmental model introduced in this study. By using the controlled [6,6’-^2^H_2_] Glc input function and measured Glx pool sizes determined with the same setup, acquisition parameters and spatial resolution, the metabolic model fully captured the time course of the measured ^2^H-Glx-C4 turnover with minimal metabolic assumptions (Figure 6), in a similar paradigm as typically performed in ^13^C-MRS labelling studies. This resulted in well determined metabolic fluxes, the free parameters of the metabolic model, as illustrated by the low standard deviation of the obtained value for Kdil and V_gt_ (*V*_*gt*_ = 0.27 ± 0.07 μmol/g/min for hippocampus and *V*_*gt*_ = 0.40 ± 0.06 μmol/g/min for striatum), in good agreement with single voxel brain values previously determined with dynamic ^13^C-MRS [25,36]. However, this study shows the additional potential of quantitative metabolic flux mapping with ^1^H-[^2^H]-FID-MRSI compared to dynamic ^13^C-MRS [22,25,26] to enable region-specific measurements, opening the way to quantitative studies of local glucose oxidative metabolism and further local pathological alterations, as illustrated here with the regional differences in Glc oxidative metabolism in the hippocampus and striatum.

The appropriateness of the metabolic model was further illustrated with MC simulations. In the field of metabolic modelling of *in vivo* data, it is critical to adapt the metabolic model to the complexity of the underlying biochemical pathways under study to properly reflect the metabolism of interest, while keeping it simple enough, based on the extent of accessible *in vivo* data, to be mathematically robust. This was shown here with the parameter recovery obtained on synthetic data with typical metabolic parameters input value of V_gt_ = 0.40 μmol/g/min, V_Lac_ = 0.21 μmol/g/min and Kdil = 0.57. Across the n = 100 MC simulations with varying noise realizations and initial conditions for the adjusted parameters, the model was able to successfully retrieve the input parameters (V_gt_ = 0.41 ± 0.06 μmol/g/min,V_Lac_ = 0.21 ± 0.10 μmol/g/min, Kdil = 0.57 ± 0.03). The calculated correlation matrix highlighted a good independence between V_gt_ and V_Lac_ retrieval, dealing with a correlation coefficient of 0.45. Regarding Kdil, its correlation coefficients relative to V_gt_ and V_Lac_ were of -0.81 and -0.52, respectively.

The study results illustrated the robustness of the metabolic model accuracy and precision in retrieving V_gt_ with varying experimental duration (Figure 7b) and noise level (Figure 7d), showing relative standard deviations (relStd) below 20% for most of the investigated conditions, and diverging significantly in the cases T_exp_ < 90 min and NF > 1.5 only. The analysis of the effect of experiment duration showed an asymptotic convergence of the adjusted model parameters towards their true values for experiment durations above 120 min (Figure 7b). Shorter experiment durations showed a trend towards an overestimation of V_gt_ and V_Lac_, in particular for experiment durations shorter than 90 min. The parameters standard deviation also increased progressively for experiment durations shorter than 150 min. For long experiment durations, a non-zero plateau is reached typically for V_gt_ and V_Lac_, illustrating the fact that these labelling fluxes are essentially characterizing the rising phase of the turnover curves, while no additional information is acquired from a longer labelling plateau measurement.

The importance of a well characterized rising phase of the ^2^H labelling curves also appears in the time resolution study (Figure 7c). The higher the number of measurement points acquired during this phase, the better the precision of the labelling fluxes V_gt_ and V_Lac_, while no major effect is observed on the accuracy of recovered value over a relatively wide range of temporal resolution. However, V_Lac_ shows a turning point around 15 min temporal resolution, above which both precision and accuracy are lost for this flux. This relates to the fact that ^2^H-Lac labelling reaches its plateau faster (Figure 7a, rising time of approximately 20 min opposite to the ^2^H-Glx 80 min) due to the small size of the Lac pool. Therefore, dynamic data sampling with Δt > 15 minutes does not enable capturing the rising phase of ^2^H-Lac turnover. Moreover, ^2^H-Lac turnover is challenging to measure for SNR reasons, due to the typically seven times smaller pool size, as compared to Glx in healthy brain tissue. On the other hand, V_gt_ regression showed good levels of accuracy and precision even in low temporal resolution conditions (Figure 7c, Δt = 20 - 30 min), still sufficiently capturing the rising phase of ^2^H-Glx turnover.

The ^2^H-Lac labelling dynamics, combined with the typical small Lac pool-size, makes V_Lac_ estimation also more sensitive to noise increases (Figure 7d, relStd > 20% for NF > 0.5, with good accuracy up to NF = 1). The model showed good accuracy in recovering V_Lac_ only for V_Lac_ < 0.4 μmol/g/min, with a progressive bias and loss in precision for higher values. Note that this applies to healthy tissue [Lac] levels (2 μmol/g) as simulated here.

The independent determination of the oxidative vs non-oxidative glucose metabolic pathways was also well illustrated in Figure 7e and Figure 7f, when separately varying V_Lac_ and V_gt_ in the input synthetic data, respectively. V_gt_ determination was strongly independent from the V_Lac_ inputs (Figure 7e, relStd < 20%), and showed a good recovery of its input value over a wide range of flux values (0.05 - 1.00 μmol/g/min, Figure 7f), underlining the reliability of the proposed metabolic modelling approach in V_gt_ determination from ^2^H-Glx turnover data. V_Lac_ was not well recovered for input V_Lac_ > 0.3 μmol/g/min, reflecting again the challenge to characterize the fast rising labelling phase of the small Lac pool with a temporal resolution of 13 min.

Future steps will involve the implementation of a new basis set for refining the Glx-C4 ^2^H-labelling characterization. In particular, a new Glu-Gln spectral pattern separation, including variations in J-coupling properties due to the presence of the ^2^H label in the molecules, could be considered. A refined basis-set could allow for the possibility of extending the ^2^H-labelling analysis to Glu and Gln separately, which would provide the characterization of the apparent Glu-Gln cycling rate, at the core of excitatory neurotransmission. A stronger magnetic field strength as 14.1T may also be exploited to favour such a labelled Glu/Gln distinguished characterization. Moreover, parallel developments towards 3D ^1^H-MRSI in the rat brain will enable acquisitions with higher SNR per voxel [37], which could translate into dynamic analysis of single voxel turnover curves. Finally, a cross-validation of the presented indirect ^2^H-MRSI measurement of glucose oxidative metabolism with direct ^2^H-MRSI acquisition protocols, focusing on Glx labelling turnover, would provide an interesting comparison for optimal SNR/spatial resolution/temporal resolution combination to achieve best accuracy and precision in TCA cycle activity determination.

## Conclusion

The proposed deuterium detection approach enabled, for the first time, the turnover characterisation of brain regional ^2^H-Glx from ^1^H-FID-MRSI following a controlled step-like [6,6’-^2^H_2_] Glc input function. The adjusted spatial and temporal resolution provided a local measurement of typical saturating exponential function for ^2^H-Glx-C4 concentration in the rat brain, characteristic for such X-nuclear labelling protocols. A dedicated compartmental metabolic model of brain glucose metabolism, adapted for the level of accessible data and for typical ^2^H label losses was developed, tested for its robustness and appropriateness, successfully describing local glucose metabolism. The combined controlled input function, metabolites pool size measurement and ^2^H turnover determination enabled the quantitative measurement of the TCA cycle metabolic flux with minimal modelling assumptions. This method opens the way to quantitative Glc oxidative flux mapping from ^1^H measurements, with no need for dedicated X-nuclei hardware.

## Acknowledgments

We acknowledge the CIBM Center for Biomedical Imaging for providing expertise and resources to conduct this study. Financial support was provided by the Swiss National Science Foundation (Projects No. 201218 and 207935) and Austrian Science Fund (WEAVE I 6037-N and P 36328-N).

## Data Availability

The data will be made available by the authors upon reasonable request.

## Compliance with Ethical Standards

This study was funded by the Swiss National Science Foundation (Grants No. 201218 and 207935) and Austrian Science Fund (Grants No. WEAVE I 6037-N and P 36328-N). All the authors declare they have no conflict of interest. All animal experiments were conducted according to federal and local ethical guidelines, and the protocols were approved by the local Committee on Animal Experimentation for the Canton de Vaud, Switzerland (VD 3892).

